# Dietary sulfur amino acid restriction improves glucose homeostasis through hepatic *de novo* serine synthesis

**DOI:** 10.1101/2025.10.16.682938

**Authors:** Andres F. Ortega, Cha Mee Vang, Ferrol I. Rome, Kaitlyn M. Andreoni, Aiden M. Phoebe, Alisa B. Nelson, Peter A. Crawford, James J. Galligan, Stanley Ching-Cheng Huang, Curtis C. Hughey

**Affiliations:** Division of Molecular Medicine, Department of Medicine, University of Minnesota, Minneapolis, MN; Department of Pharmacology and Toxicology, University of Arizona, Tucson, AZ; Department of Biochemistry, Molecular Biology, and Biophysics, University of Minnesota, Minneapolis, MN; Pelotonia Institute for Immuno-oncology, Comprehensive Cancer Center – James Cancer Hospital and Solove Research Institute, The Ohio State University, Columbus, OH; Department of Microbial Infection and Immunity, College of Medicine, The Ohio State University, Columbus, OH

**Keywords:** Sulfur amino acid restriction, De novo serine synthesis, Glucose homeostasis, Liver intermediary metabolism

## Abstract

Dietary sulfur amino acid restriction (SAAR) improves whole-body glucose homeostasis, elevates liver insulin action, and lowers liver triglycerides. These adaptations are associated with an increased expression of hepatic *de novo* serine synthesis enzymes, phosphoglycerate dehydrogenase (PHGDH) and phosphoserine aminotransferase 1 (PSAT1). This study tested the hypothesis that enhanced hepatic serine synthesis is necessary for glucose and lipid adaptations to SAAR. Hepatocyte-specific PSAT1 knockout (KO) mice and wild type (WT) littermates were fed a high-fat control or SAAR diet. In WT mice, SAAR increased liver PSAT1 protein (∼70-fold), serine concentration (∼2-fold), and ^13^C-serine (∼20-fold) following an intravenous infusion of [U-^13^C]glucose. The elevated liver serine and partitioning of circulating glucose to liver serine by SAAR were attenuated in KO mice. This was accompanied by a blunted improvement in glucose tolerance in KO mice fed a SAAR diet. Interestingly, SAAR decreased liver lysine lactoylation, a SAA-supported post-translational modification known to inhibit PHGDH enzymatic activity. This suggests dietary SAAR may increase serine synthesis, in part, by lowering lysine lactoylation. Beyond glucose metabolism, dietary SAAR reduced body weight, adiposity, and liver triglycerides similarly in WT and KO mice. Collectively, these results demonstrate that hepatic PSAT1 is necessary for glucose, but not lipid, adaptations to SAAR.

**Graphical Abstract:** Schematic representation of liver glucose adaptations to SAAR

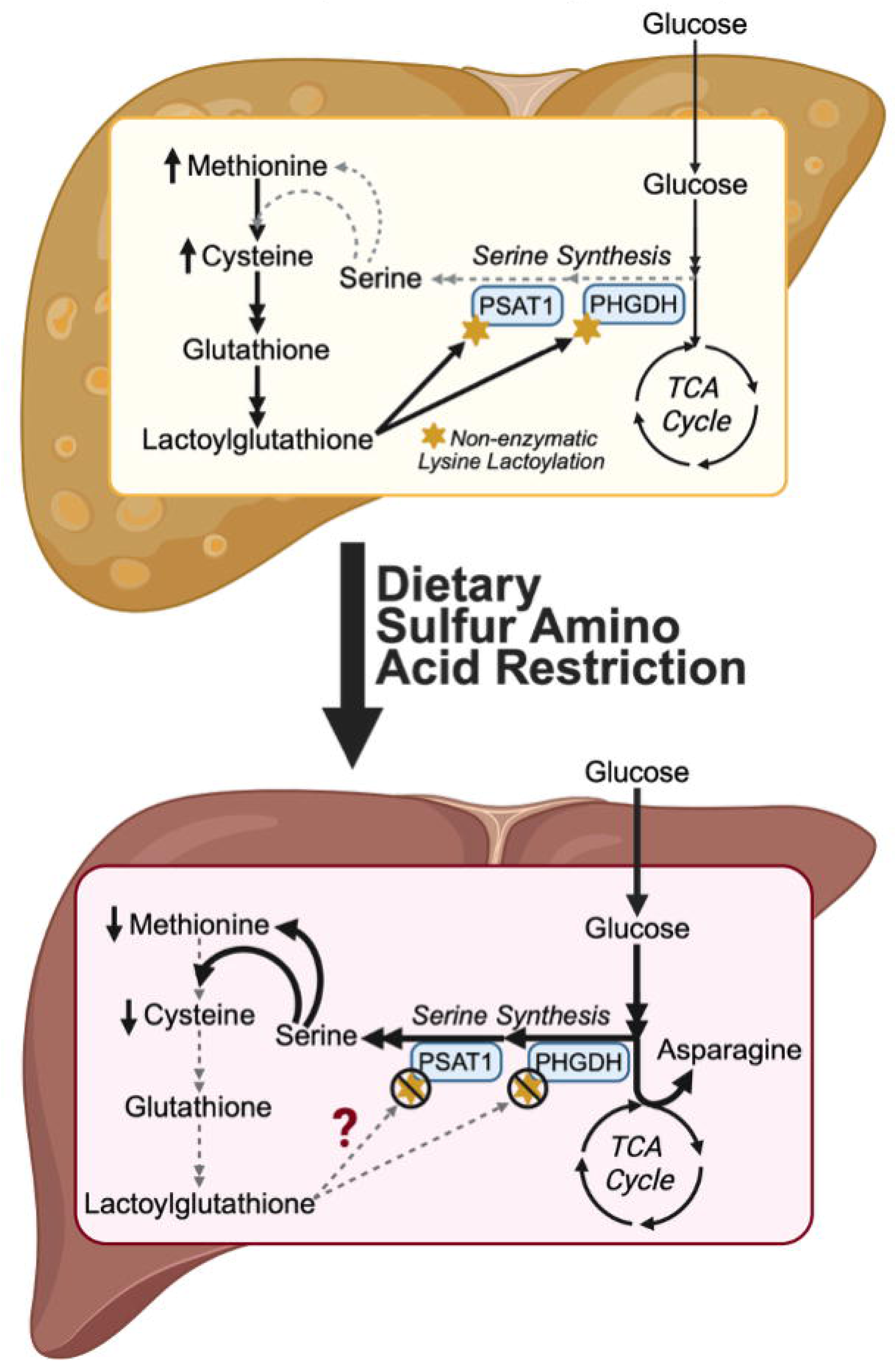

## 1. INTRODUCTION

Dietary sulfur amino acid restriction (SAAR) provokes metabolic adaptations that are advantageous in preventing and treating obesity and its comorbidities [1]. In rodents, SAAR enhances glucose and insulin tolerance [2–7]. The use of hyperinsulinemic-euglycemic clamps combined with isotope tracers revealed that the increased insulin action is due to a greater suppression of endogenous glucose production and heightened glucose disposal by skeletal muscle, heart, and adipose tissue [8–11]. Concurrently, SAAR decreases adiposity and liver triglycerides [1–4]. It is important to note that these metabolic responses to SAAR are not confined to rodents. Short-term dietary SAAR in humans elicits decreases in body weight, body mass index (BMI), and liver fat [12–14]. The SAAR-mediated adaptations are partly due to a liver-derived increase in fibroblast growth factor 21 (FGF21) [1]. The transcriptional activation of hepatic FGF21 increases whole-body energy expenditure, which contributes to reduced adiposity by SAAR [9]. FGF21 is necessary for the enhanced insulin-stimulated cardiac glucose uptake [10; 11]. In contrast, SAAR lowers liver triglycerides similarly in wild type (WT) and FGF21-deficient mice[9]. Prior work indicates the enhanced liver insulin signaling by SAAR is also independent of FGF21 [1; 11]. Together, these findings suggest that the increased FGF21 mediates several extrahepatic responses to SAAR, but may not be critical for the hepatic responses. Given this, the mechanisms underlying the liver glucose and lipid adaptations to SAAR, and the extent to which these responses are hepatocyte-dependent, remain to be clearly defined.

Elevated expression of liver *de novo* serine synthesis pathway enzymes, phosphoglycerate dehydrogenase (PHGDH) and phosphoserine aminotransferase 1 (PSAT1), is among the most reproducible responses to dietary SAAR, making their increase a marker of a successful dietary intervention [2; 9; 10; 15–23]. *De novo* serine synthesis is sourced from 3-phosphoglycerate, an intermediate of both glycolysis and gluconeogenesis [24]. Thus, a SAAR-mediated increase in hepatic serine synthesis could promote greater glucose disposal and/or divert gluconeogenic flux away from glucose production. Liver-specific PHGDH knockout (KO) mice fed a chow diet show higher adiposity and reduced glucose tolerance [25]. In agreement with PHGDH KO mice, PSAT1 knockdown impairs glucose and insulin tolerance [26]. Conversely, PSAT1 overexpression enhances glucose and insulin tolerance [26]. Beyond glucose homeostasis, the overexpression of PHGDH has been shown to lower liver lipids [27]. These prior results support that PHGDH and PSAT1 action impacts whole-body and liver-specific glucose and lipid homeostasis.

The experiments presented herein tested the hypothesis that hepatic PSAT1 is required for the liver glucose and lipid adaptations to SAAR to be fully realized. Liver-specific PSAT1 KO mice and WT littermates were fed a high-fat control or SAAR diet. Comprehensive phenotyping approaches were completed. These included glucose and insulin tolerance tests to define whole-body glucose homeostasis, ^13^C-isotope tracing in conscious unrestrained mice to determine the tissue-specific fate of glucose and serine *in vivo*, and metabolomics to quantify metabolite concentrations. Immunoblotting and proteomics were used to identify molecular drivers of liver metabolism. The results of this study show that hepatic PSAT1 is necessary for whole-body and liver-specific glucose, but not lipid, adaptations to dietary SAAR.

## 2. MATERIALS AND METHODS

### 2.1. Mouse models and husbandry

All procedures using mice were approved by the University of Minnesota Institutional Animal Care and Use Committee. PSAT1 floxed mice on a C57BL/6N background [28] were backcrossed onto a C57BL/6J background for 5 generations. Hepatic-specific PSAT1 KO were then generated by breeding mice expressing Cre recombinase under control of the albumin promoter (B6.Cg-Tg(Alb-cre)21Mgn/J) with PSAT1 floxed mice on a C57BL/6J background. Floxed PSAT1 littermates lacking expression of Cre recombinase were used as WT controls. C57BL/6J mice were used for select studies. Mice were maintained on a chow diet (Teklad Global 18% Protein Rodent Diet 2918, Madison, WI) from weaning at three weeks of age until six weeks of age. At six weeks of age mice received one of three diets (unless otherwise stated): low-fat SAA control (0.85% kcal from methionine; 10% kcal from fat; LF-CTRL; A11051302B, Research Diets Inc., New Brunswick, NJ), high-fat SAA control (0.85% kcal from methionine; 60% kcal from fat; HF-CTRL; A11051306, Research Diets Inc., New Brunswick, NJ), and HF-SAAR (0.125% kcal from methionine; 60% kcal from fat; A11051305, Research Diets Inc., New Brunswick, NJ). The full dietary composition for these three diets is detailed in Supplementary Table 1. Food and water were provided ad libitum. Mice were housed with cellulose bedding (Cellu-nest™, Shepherd Specialty Papers, Watertown, TN) in temperature-and humidity-controlled conditions maintained on a university instituted 14:10 hour light:dark cycle. At 12 weeks of age (6 weeks on diet), mice underwent metabolic testing and were euthanized via cervical dislocation. Tissues were rapidly excised, freeze-clamped in liquid nitrogen, and stored at-80°C. One cohort of mice was euthanized at 26 weeks of age (20 weeks on diet) using the same procedures described above. All mice used in this study were males.

### 2.2. Body Composition

Body composition was measured in mice fed ad libitum at 10-11 weeks of age using an EchoMRI-100^TM^ Body Composition Analyzer (EchoMRI LLC, Houston, TX).

### 2.3 Glucose and insulin tolerance tests

For glucose tolerance tests (GTT), mice were fasted for five hours and injected intraperitoneally (i.p.) with dextrose (Hospira, Inc., Lake Forest, IL) at a dose of 2 g·kg^-1^ lean weight. For insulin tolerance tests (ITT), mice were fasted for five hours and received insulin (Humulin R human, Eli Lilly and Company, Indianapolis, IN) i.p. at a dose of 0.75 U·kg^-1^ lean weight. During both procedures, tail-blood glucose was measured at 0, 15, 30, 45, 60, 90, and 120 minutes with a Contour blood glucose meter (Ascensia Diabetes Care, Parsippany, NJ). A cohort of mice had blood samples obtained at 0 and 15 minutes after the insulin administration for plasma insulin concentrations and were euthanized immediately after for assessing indices of insulin signaling in tissues.

### 2.4. Insulin pellet implantation

At 7 weeks of age, C57BL/6J mice had one sustained-release insulin pellet implanted subcutaneously (∼0.1 U/day per implant; LinBit; LinShin Canada, Toronto, ON) according to manufacturer guidelines. Briefly, under isoflurane anesthesia, a 1 cm diameter area of hair was clipped and disinfected. A small incision in the skin was made with a 16-gauge needle, and a sterile 12-gauge trocar was used to deliver the pellet subcutaneously. Procedures in control mice were identical, except the skin incisions were not performed. Mice were group housed following the procedure.

### 2.5. Vascular catheter surgeries

At 11 weeks of age, C57BL/6J mice had catheters were surgically implanted in the jugular vein and carotid artery for isotope infusion and sampling protocols as previously described [21; 29]. The exteriorized ends of the implanted catheters were flushed with 200 U·mL^-1^ heparinized saline and sealed with stainless-steel plugs. Following surgery, mice were housed individually and provided 5-7 days of post-operative recovery prior to stable isotope infusion studies. Liver-specific PSAT1 KO and WT littermates underwent identical surgical procedures with the exception that only the jugular vein catheter was implanted. All mice were within 10% of pre-surgical weight prior to stable isotope infusions.

### 2.6. Stable-isotope tracing: [U-^13^C]glucose intravenous infusion in the conscious, unrestrained mouse

During the first hour of the housing light cycle, both food and water were withdrawn for the duration of the experiment (5.5 hours). Four hours into the fast, the exteriorized vascular catheters were connected to infusion syringes. Following a one-hour acclimation, a 60 mg·kg^-1^·min^-1^ continuous intravenous infusion of [U-^13^C]glucose was performed for 30 minutes. C57BL/6J mice had arterial samples obtained for measuring blood glucose prior to and every 10 minutes during the [U-^13^C]glucose infusion to ensure the infusion rate was sufficient to achieve glucose concentrations comparable to those observed during a GTT. Mice were euthanized and tissue collected immediately following the [U-^13^C]glucose infusion.

### 2.7. Stable-isotope tracing: In-vivo [U-^13^C]serine intraperitoneal injection

Food and water were withdrawn for 5 hours starting within the first hour of the light cycle. [U-^13^C]serine was administered i.p. at a dose of 100 mg·kg^-1^. Fifteen minutes following the i.p. injection, mice were euthanized and tissues harvested.

### 2.8. Hormone and metabolite analysis

Plasma insulin was determined using the Mercodia Ultrasensitive Mouse Insulin ELISA (Mercodia AB, Winston Salem, NC). Liver triglycerides were measured with the Triglycerides - Liquid Reagent Set (Pointe Scientific, Inc., Lincoln Park, MI). Liver glycogen was quantified as previously described [21; 30]. All other metabolites were measured using untargeted metabolomics as detailed below. Tissue metabolite levels were quantified from mice that were fasted 7 hours.

### 2.9. Untargeted metabolomics and ^13^C-isotope tracing untargeted metabolomics (^13^C-ITUM) pipeline

Extraction and analytical methods were completed as previously described [31–33] with minor modifications. Briefly, lyophilized liver samples (∼2 mg) were homogenized in 1000 µL of 2:2:1 (v/v/v) acetonitrile (ACN):methanol (MeOH):water and subjected to three (30 second) cycles of vortexing, one minute of flash-freezing in liquid nitrogen, and 10 minutes of sonication at 25°C. Homogenized samples were stored at −20°C for 1 hour and then centrifugated at 15,000 × g for 10 minutes at 4 °C. Subsequently, the supernatant was transferred to a fresh tube and dried by SpeedVac overnight. Liver extracts were reconstituted in 100 µL of 1:1 (v/v) ACN:water, sonicated for two (five minute) cycles, vortexed for 1 minute, and then stored at 4°C for one hour. Finally, samples were centrifuged at 15,000 × g at 4 °C for 10 minutes and the supernatants transferred to LC-MS vials. Metabolites were separated using a Thermo Fisher Scientific Vanquish UHPLC system and an Atlantis Premier UPLC BEH Z-HILIC column (186009979, Milford, MA). Mobile Phase A was 90% water with 15 mM ammonium bicarbonate and Mobile Phase B was 90% ACN with 15 mM ammonium bicarbonate. The total run time was 10 minutes, flow rate was 0.5 mL·min^-1^, and column chamber temperature was 30°C. Separated metabolites were detected using a Thermo Fisher Scientific QExactive Plus hybrid quadrupole-orbitrap mass spectrometer, fitted with a heated electrospray ionization source operated in negative and positive polarity mode. Two sample types were run to aid in chemical feature identification: 1) a standard mix consisting of authentic standards for all expected analytes and 2) a pooled sample containing only the naturally occurring samples, analyzed via data-dependent analysis (DDA)–tandem mass spectrometry (MS/MS), using IE omics script and R-Studio [34]. Data processing and initial analysis were performed using Thermo Compound Discoverer 3.3. Raw mass spectra were uploaded, background ions were removed, retention times (RT) for detected signals were aligned across samples, and chemical formulas were predicted. Grouped chemical features were profiled to determine compound identity based on: 1) the *m*/*z* predicted from the chemical formula, 2) the RT compared to an authentic external standard, and 3) the MS/MS fragmentation pattern, compared to in-house standards and online databases. For ITUM, putatively identified metabolites were carried forward and subjected to ^13^C-isotope enrichment with correction for natural abundance. To carry out ^13^C-isotope tracing, all [^13^C] mass isotopomers within the isotopic envelope of each identified metabolite were determined based on the diagnostic shift in *m*/*z* (Δ*m*/*z* = 1.0033 Da, natural abundance, 1.11% of all carbon) induced by the presence of ^13^C-labeled compounds. Raw ion counts for each isotopologue were plotted against RT to generate extracted ion chromatograms, and then the area under the curve for each isotopologue was extracted, summed, and expressed as a percentage of the total pool. After natural abundance correction, the fractional intensities were then graphed as a function of ^13^C-carbon content, generating mass isotopologue distributions for each detected metabolite. Fractional enrichment reports on the percentage of molecules in a metabolite pool that are ^13^C-enriched and was calculated by the summation of the relative intensities (% of the total pool) for all ^13^C mass isotopologues detected. Untargeted metabolomics to determine relative metabolite pools sizes was performed in livers from mice fasted 7 hours to avoid the potential impact of ^13^C-isotope administration on metabolite abundance.

### 2.10. Targeted analysis of glyoxalase cycle intermediates

Reduced glutathione (GSH), lactoylglutathione (LGSH), and hemithioacetal (HTA) were quantified in liver as previously described [35]. Briefly, samples were prepared in a sulfosalicylic acid solution containing the internal standard, GSH-(Gly-^13^C_2_,^15^N). The protein precipitate was removed via centrifugation and supernatants chromatographed using a Shimadzu LC system equipped with an Atlantis C_18_ column (186001291; Waters, Milford, MA). Multiple reaction monitoring-mass spectrometry (MRM-MS) was performed in positive ion mode using an AB SCIEX 6500+ QTRAP.

### 2.11. Quantitative analysis of arginine and lysine modifications

Lysine lactoylation (lactoyllys), lysine acetylation (acetyllys), methylglyoxal-derived hydroimidazolone 1 (MG-H1), and carboxyethylarginine (CEA) were determined as previously described [35; 36]. Liver protein was precipitated and resuspended in ammonium bicarbonate containing internal standards. Samples were then digested to single amino acids, acidified with heptofluorobutyric acid, chromatographed using a Shimadzu LC system equipped with an Eclipse XDB-C8 column (930990-906; Agilent, Santa Clara, CA), and analyzed by MRM-MS. The post-translational modifications (PTMs) were normalized to Leu to control for digestion [36; 37].

### 2.12. Immunoblotting

Tissue homogenates were prepared as previously detailed [21; 29]. Tissue proteins (15 µg) were separated by electrophoresis on a NuPAGE 4-12% Bis-Tris gel (Invitrogen, Carlsbad, CA, USA) and transferred to a PVDF membrane. Immunoblotting was performed with the primary antibodies and secondary antibodies listed in Supplementary Table 2. The PVDF membranes were treated with a chemiluminescent substrate (Thermo Fisher Scientific, Waltham, MA, USA) and imaged using a ChemiDoc Imaging system and Image Lab software (Bio-Rad, Hercules, CA, USA). Total protein was assessed via BLOT-FastStain (G-Bioscience, St. Louis, MO, USA) and used as the loading control. Densitometry was evaluated using ImageJ software.

### 2.13. Statistical Analysis

GraphPad Prism software (GraphPad Software LLC., San Diego, CA) was used to perform Student’s t-test, two-way ANOVAs followed by Tukey’s post-hoc tests, and repeated measures ANOVAs followed by Šidák’s post-hoc tests as appropriate to detect statistical differences (p<0.05). All data are reported as mean ± SEM. Data points greater than two standard deviations from the mean were considered outliers and excluded from analysis.

## 3. RESULTS

### 3.1. Elevated glucose partitioning to liver serine by SAAR is linked to improved glucose tolerance

In C57BL/6J mice, 6 weeks of dietary intervention showed a greater gain in body weight, higher adiposity, and similar lean mass in the HF-CTRL compared to LF-CTRL group (Fig. 1A and B). Mice fed a HF-SAAR diet had a lower body weight, decreased fat mass, and increased lean mass percentage relative to both the LF-CTRL and HF-CTRL mice (Fig. 1B). A HF-CTRL diet elevated liver triglycerides compared to mice fed the LF-CTRL diet, while the HF-SAAR diet prevented the increase in liver triglycerides and lowered liver weight (Fig. 1C). Dietary SAAR also lowered liver glycogen and fasting blood glucose (Fig. 1D and E). HF-CTRL mice showed impaired glucose tolerance, but not insulin tolerance, compared to LF-CTRL mice (Fig. 1F and G). Conversely, glucose and insulin tolerance were improved in mice fed the HF-SAAR diet relative to mice fed the LF-CTRL and HF-CTRL diets (Fig. 1F and G). Plasma insulin was significantly reduced in HF-SAAR mice (Fig 1H). This is notable because chronically elevated insulin has been proposed to promoted gain in adiposity and dysregulated glucose control [38–40]. To test if the lowered circulating insulin levels in HF-SAAR mice was contributing to the metabolic adaptations to SAAR, a subcutaneous, sustained-release insulin pellet was implanted in HF-SAAR mice to raise insulin. The persistent delivery of exogenous insulin from the subcutaneous pellet over 5 weeks did not impact the glucose and lipid adaptations to SAAR (Suppl. Fig. 1). This suggests that the mechanisms underlying SAAR are independent of circulating insulin concentrations.

**Figure 1.**
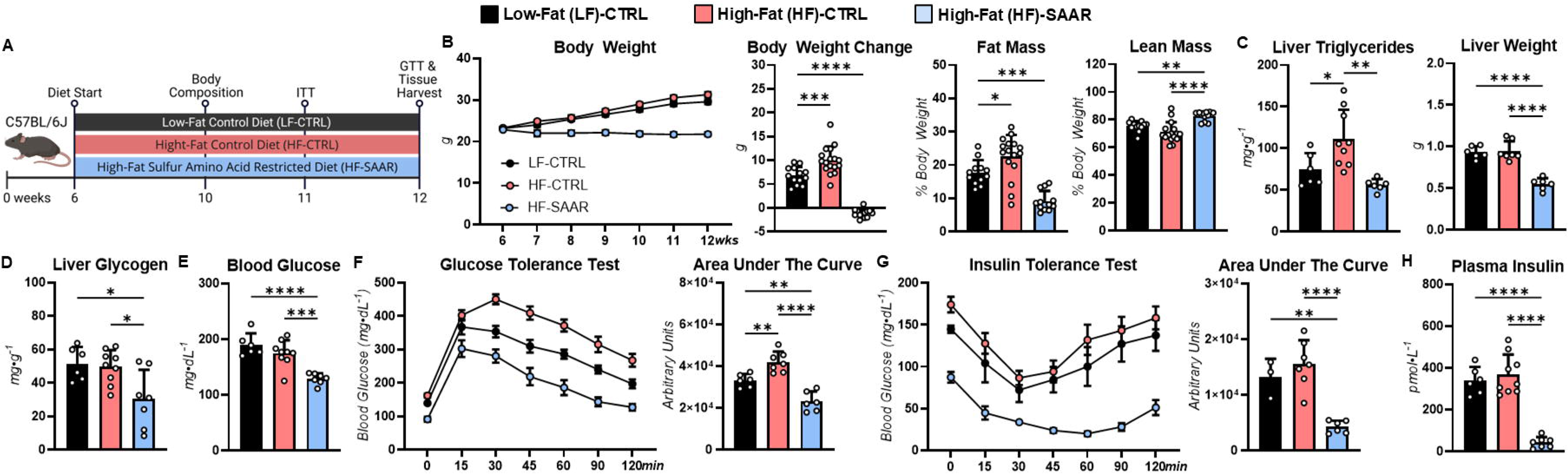
Sulfur amino acid restriction (SAAR) prevents obesity and improves systemic glucose control in C57BL/6J mice. **A.** A schematic representation of the study design. C57BL/6J male mice were fed a low-fat control (LF-CTRL), high-fat control (HF-CTRL), or HF-SAAR diet for 6 weeks starting at 6 weeks of age. **B.** Body weight time course (g), body weight change over the 6 weeks on diet (g), fat mass (% of body weight), and lean mass (% of body weight). **C.** Liver triglycerides (mg·g^-1^) and liver weight (g). **D.** Liver glycogen (mg·g^-1^)**. E.** Blood glucose (mg·dL^-1^). **F.** Glucose excursion and area under the curve (AUC) during a glucose tolerance test. **G.** Glucose excursion and AUC during an insulin tolerance test. **H.** Plasma insulin (pmol·L^-1^). n=3-17. *p<0.05, **p<0.01, ***p<0.001, ****p<0.0001.

Untargeted metabolomics revealed that liver cysteine was higher in HF-CTRL compared to LF-CTRL mice (Fig. 2A and 2B). As expected, liver cysteine and taurine were decreased ∼5-fold in mice fed the HF-SAAR diet (Fig. 2B). The HF-SAAR diet led to a markedly increased in liver PHGDH (∼5-fold) and PSAT1 (∼70-fold) protein expression (Fig. 2C). The elevated expression of serine synthesis enzymes was associated with higher liver serine and glycine levels in HF-SAAR relative to both CTRL diet groups (Fig. 2E). The profound increase in PSAT1 protein in the HF-SAAR group made it difficult to resolve differences in PSAT1 in the livers of mice on the LF-and HF-CTRL diets. However, immunoblots performed specifically to compare the LF-CTRL and HF-CTRL groups showed that high-fat feeding decreased liver PSAT1 (Fig. 2D). This was accompanied lower serine concentrations in LF-CTRL versus HF-CTRL mice (Fig. 2E). To test if the increased liver serine and glycine were being sourced from circulating glucose, [U-^13^C]glucose was intravenously infused in conscious, unrestrained mice fed a LF-CTRL or a LF-SAAR diet for 6 weeks (Fig. 2F). ^13^C-labeling of liver serine and glycine was significantly higher in LF-SAAR mice compared to LF-CTRL, with no alterations observed in labeled glucose (Fig. 2H). Collectively, our data show that dietary SAAR improves glucose and insulin tolerance, which is associated with increased partitioning of circulating glucose to liver serine synthesis.

**Figure 2.**
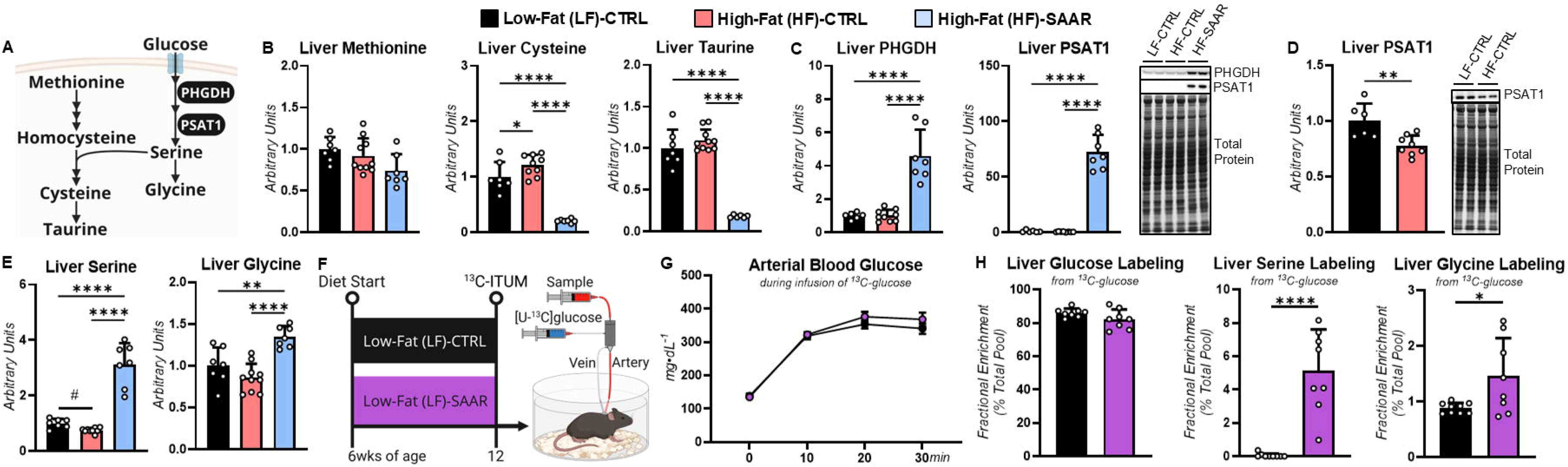
Sulfur amino acid restriction (SAAR) increases liver glucose partitioning to *de novo* serine synthesis in C57BL/6J mice. **A.** A schematic representation of the select reactions involved in *de novo* serine synthesis and SAA synthesis and catabolism. **B.** Liver methionine, cysteine, and taurine (arbitrary units) in C57BL/6J male mice fed a low-fat control (LF-CTRL), high-fat control (HF-CTRL), or HF-SAAR diet for 6 weeks starting at 6 weeks of age. **C.** Liver phosphoglycerate dehydrogenase (PHGDH) and phosphoserine aminotransferase 1 (PSAT1) as determined by immunoblotting and representative immunoblots in mice fed a LF-CTRL, HF-CTRL, and HF-SAAR diet. **D.** Liver PSAT1 as determined by immunoblotting and representative immunoblots in mice fed a LF-CTRL and HF-CTRL diet only. **E.** Liver serine and glycine (arbitrary units). **F.** A schematic representation of the study design. C57BL/6J mice were fed a LF-CTRL (0.88% kcal from methionine, 510072, Dyets Inc., Bethlehem, PA) or low-fat SAAR (LF-SAAR; 0.18% kcal from methionine, 510071, Dyets Inc., Bethlehem, PA) for 6 weeks starting at 6 weeks of age. See Supplementary Table 3 for diet details. A 30-minute intravenous infusion of [U-^13^C]glucose in the conscious, unrestrained mouse was completed at 12 weeks of age. **G.** Arterial blood glucose (mg·dL^-1^). **H.** Fractional enrichment (% total pool) of liver glucose, serine and glycine from [U-^13^C]glucose. n=6-10. ^#^p<0.05 by *t*-test, *p<0.05, **p<0.01, ***p<0.001, ****p<0.0001 by ANOVA.

### 3.2. Liver PSAT1 is required for SAAR-induced improvements in systemic glucose homeostasis

To assess whether hepatic serine synthesis is essential for the metabolic benefits of dietary SAAR on glucose control, we studied liver-specific PSAT1 KO mice and WT littermates fed a HF-CTRL or-SAAR diet (Fig. 3A). The HF-SAAR diet prevented a gain in body weight and adiposity in both genotypes (Fig. 3B and C). When fed a HF-CTRL diet, WT and liver PSAT1 KO mice showed similar glucose tolerance (Fig. 3D). This was expected given that high-fat feeding lowers liver PSAT1 in WT mice (Fig. 2D), a phenotype closely resembling liver PSAT1 KO mice. On a HF-SAAR diet, liver-specific PSAT1 KO mice showed a greater excursion of glucose at the 15-and 30-minute time points during the glucose tolerance test compared to WT mice, although the area under the curve was comparable between genotypes (Fig. 3D). Loss of hepatic PSAT1 in mice fed HF-CTRL diet delayed the normalization of circulating glucose during an insulin tolerance test relative to WT littermates (Fig. 3E). Six weeks of HF-SAAR feeding improved insulin tolerance similarly in WT and KO mice compared to HF-CTRL groups (Fig. 3E). Concordant with the C57BL/6J findings (Fig. 1), HF-SAAR feeding lowered plasma insulin levels relative to mice fed a HF-CTRL diet in both genotypes (Fig. 3F). Fifteen minutes after an insulin injection, the liver p-Akt-to-Akt ratio was higher in both genotypes fed the HF-SAAR diet (Fig. 3G). This suggests liver insulin signaling was enhanced similarly in WT and KO mice following SAAR. Dietary SAAR lowered liver glycogen in WT and liver PSAT1 KO mice (Fig. 3H). However, this was independent of changes key enzymes regulating glycogen synthesis and breakdown. Specifically, total protein and the phospho-to-total protein ratio for glycogen synthase (GS) and glycogen phosphorylase (PYGL) were similar between groups (Fig. 3I). Together, these data indicate that liver PSAT1 is necessary for dietary SAAR to improve whole-body glucose tolerance.

**Figure 3.**
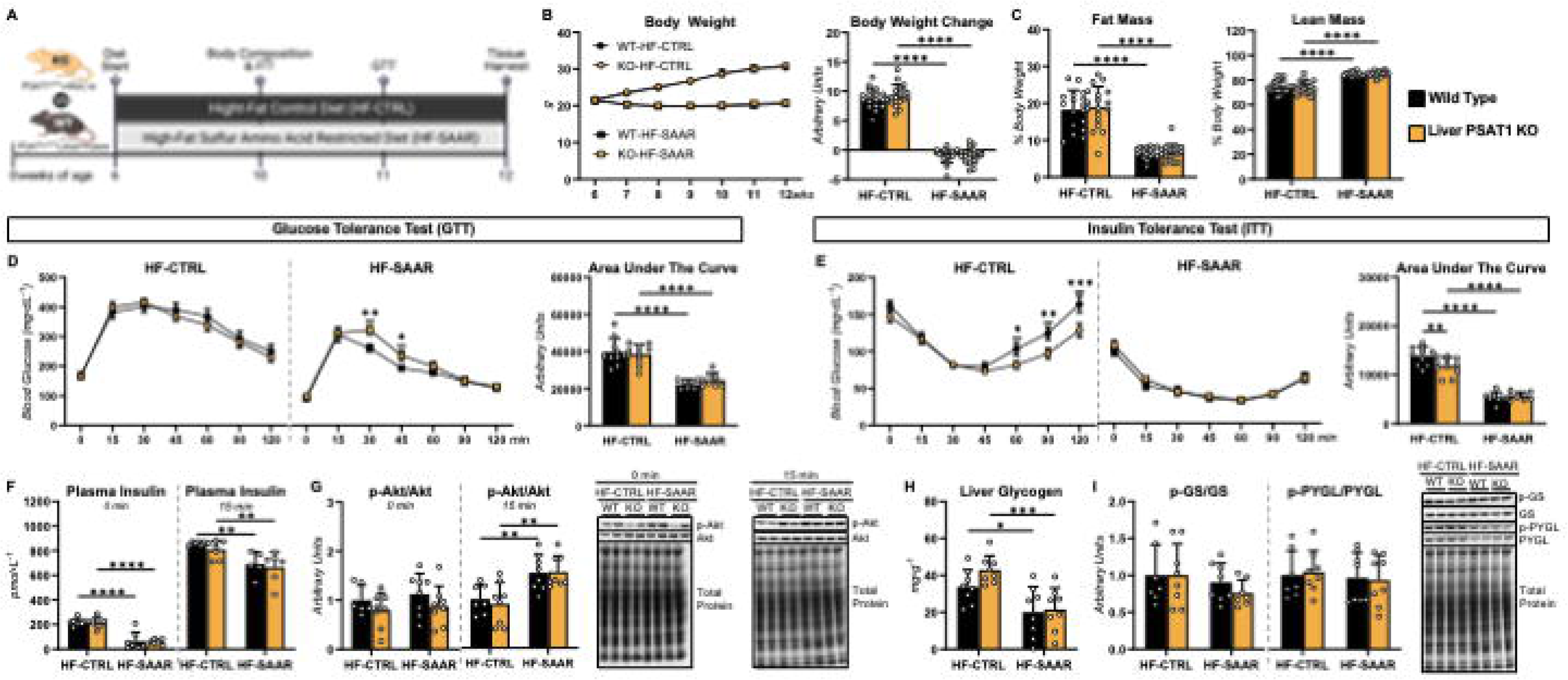
Liver PSAT1 is required for systemic glucose adaptations to sulfur amino acid restriction (SAAR). **A.** A schematic representation of the study design. Liver-specific PSAT1 knockout (KO) mice and wild type (WT) littermates were fed a high-fat control (HF-CTRL) or HF-SAAR diet for 6 weeks. **B.** Body weight time course (g) and body weight change over 6 weeks on diet (g). **C.** Fat mass and lean mass (% of body weight). **D.** Glucose excursion curves and area under the curve (AUC) for glucose tolerance tests. **E.** Glucose excursion curves and AUC for insulin tolerance tests. **F.** Plasma insulin (pmol·L^-1^) before (0 min) and 15 minutes after an insulin injection. **G.** Liver phosphor-Akt (p-Akt)-to-Akt ratio as determined by immunoblotting and representative immunoblots. **H.** Liver glycogen (mg·g^-1^). **I.** Liver phospho-glycogen synthase (p-GS)-to-GS ratio and phospho-glycogen phosphorylase (p-PYGL)-to-PYGL ratio as determined by immunoblotting and representative immunoblots. n=5-18. *p<0.05, **p<0.01, ***p<0.001, ****p<0.0001.

### 3.3. Increased glucose partitioning to serine following SAAR is abolished by loss of hepatic PSAT1

Next, we aimed to further connect the attenuated improvement in glucose tolerance observed in liver PSAT1 KO mice fed a HF-SAAR diet to inability of hepatocytes to synthesize serine from glucose (Fig. 4A). The HF-SAAR diet elevated serine and glycine synthesis enzymes, PHGDH, PSAT1 (WT mice only), and serine hydroxymethyltransferase 2 (SHMT2), in the livers of mice (Fig. 4B). Under post-absorptive conditions, liver glucose, serine, and glycine were comparable between WT and KO mice fed a HF-CTRL diet (Fig. 4C). Dietary SAAR lowered liver glucose in WT, but not liver PSAT1 KO mice (Fig. 4C). Liver serine and glycine were higher in WT mice fed a HF-SAAR diet compared to WT mice on a HF-CTRL diet (Fig. 4C); however, the increase in liver serine and glycine by SAAR was blunted in liver PSAT1 KO mice (Fig. 4C). Consistent with these metabolite levels, a HF-SAAR diet prompted a rise in the ^13^C-labeling of liver serine and glycine in WT mice intravenously infused with [U-^13^C]glucose (Fig. 4D). Loss of liver PSAT1 attenuated the increased in serine and glycine ^13^C-labeling by SAAR (Fig. 4D). These data imply that the improved glucose tolerance by dietary SAAR is associated with enhanced partitioning of circulating glucose to liver serine synthesis.

**Figure 4.**
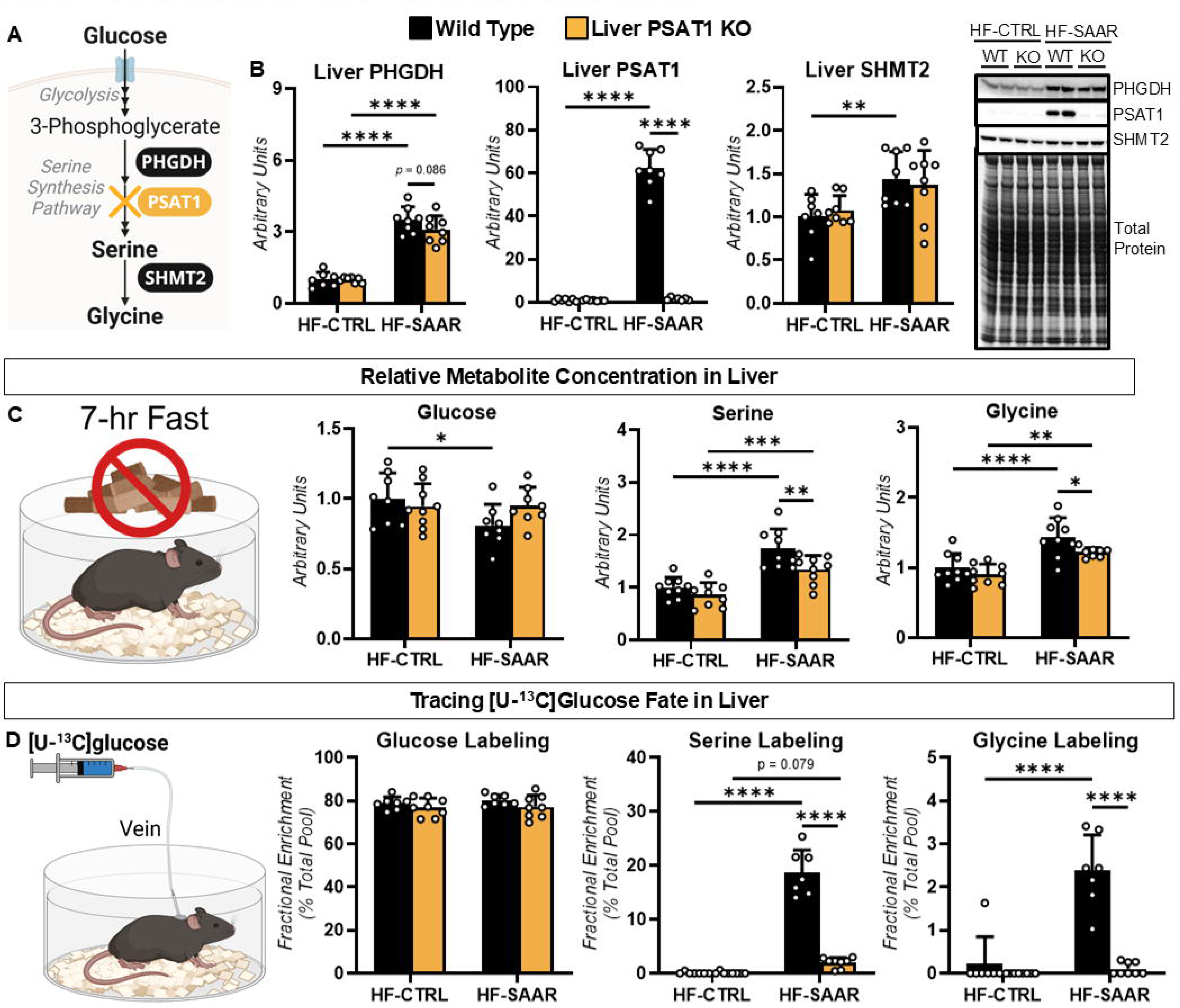
Sulfur amino acid restriction (SAAR) increases liver *de novo* serine synthesis from glucose. **A.** A schematic representation of select metabolites and enzymes involved in *de novo* serine synthesis from glucose. **B.** Liver phosphoglycerate dehydrogenase (PHGDH), phosphoserine aminotransferase 1 (PSAT1), and serine hydroxymethyltransferase 2 (SHMT2) as determined by immunoblotting and representative immunoblots in wild type (WT) and liver-specific PSAT1 knockout (KO) mice fed a high-fat control (HF-CTRL) or HF-SAAR diet for 6 weeks. **C.** Liver glucose, serine, and glycine (arbitrary units) in 7-hour fasted mice. **D.** Fractional enrichment (% of total pool) of liver glucose, serine, and glycine following a 30-minute intravenous infusion of [U-^13^C]glucose. n=7-9. *p<0.05, **p<0.01, ***p<0.001, ****p<0.0001.

### 3.4. Serine sourced from glucose following SAAR is preferentially fated to glutathione

Serine is a precursor for cysteine synthesis via transsulfuration and donates a one-carbon units to support methionine regeneration within the methionine cycle [41]. Given this, one could speculate that the elevated diversion of glucose to serine is an adaptive response to prevent limitations in SAA availability that may arise from dietary SAAR. To further test this hypothesis, liver proteins, metabolite concentrations, and metabolite labeling from [U-^13^C]glucose and [U-^13^C]serine administration were determined (Fig. 5). Regardless of diet, loss of hepatic PSAT1 did not impact proteins involved in transsulfuration or cysteine catabolism (Fig. 5A and 5B). Dietary SAAR increased cystathionine γ-lyase (CSE; involved in cysteine production), decreased cysteine dioxygenase 1 (CDO1; involved in cysteine catabolism to taurine), and elevated GSH synthesis enzymes, glutamate-cysteine ligase catalytic subunit (GCLC) and glutathione synthetase (GSS), in both genotypes (Fig. 5A and 5B). SAAs and related metabolites were comparable between WT and liver PSAT1 KO mice fed a HF-CTRL diet (Fig. 5C). The HF-SAAR diet decreased liver cysteine, taurine, GSH, and oxidized glutathione (GSSG) in both genotypes compared to their counterparts fed the HF-CTRL diet (Fig. 5C). Consistent with metabolite levels, ^13^C-labeling of liver SAA and related metabolites following a 30-minute [U-^13^C]glucose infusion was similar between WT and Liver PSAT1 KO mice on a HF-CTRL diet (Fig. 5D). HF-SAAR feeding increased ^13^C-labeling from glucose in liver methionine in both WT and liver PSAT1 KO mice (Fig. 5D). Fractional enrichment in liver cysteine was not detectable due to the low cysteine pool in mice fed a HF-SAAR diet; however, downstream fates of cysteine were measured. The [U-^13^C]glucose infusion showed lower ^13^C-taurine, and increased ^13^C-GSH and ^13^C-GSSG in the livers of both WT and KO mice fed the HF-SAAR diet relative to the HF-CTRL mice (Fig. 5D), with the SAAR-mediated increase in GSH and GSSG ^13^C-labeling from [U-^13^C]glucose being blunted in liver PSAT1 KO mice compared to their WT littermates (Fig. 5D). To more directly assess the impact of dietary SAAR on serine fate, [U-^13^C]serine was administered to WT mice fed a HF-CTRL or a HF-SAAR diet for 6 weeks (Fig. 5E). ^13^C-labeling was increased in liver methionine, taurine, GSH, and GSSG in the HF-SAAR mice compared to the HF-CTRL (Fig. 5E). The rise in ^13^C-labeling was particularly pronounced for GSH (∼12-fold) and GSSG (∼37-fold) (Fig. 5E). Together, these results suggest that serine synthesized *de novo* from glucose is important for supporting GSH provision during SAAR.

**Figure 5.**
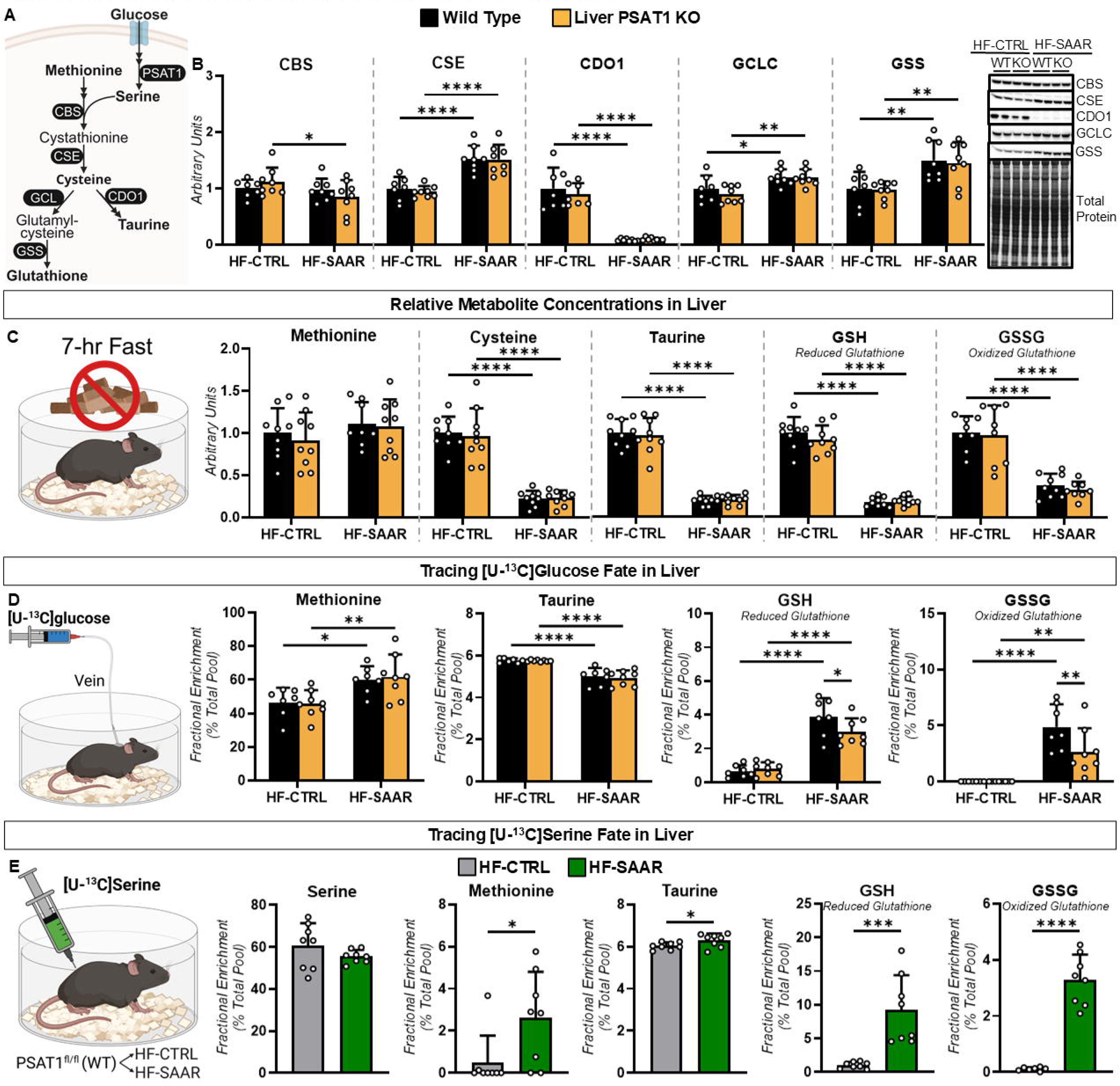
Sulfur amino acid restriction (SAAR) increases partitioning of glucose-sourced serine to liver glutathione. **A.** A schematic representation of the select reactions involved in *de novo* serine synthesis and SAA synthesis and catabolism. **B.** Liver cystathionine β-synthase (CBS), cystathionine γ-lyase (CSE), cysteine dioxygenase 1 (CDO1), glutamate-cysteine ligase catalytic subunit (GCLC), and glutathione synthetase (GSS) as determined by immunoblotting and representative immunoblots in wild type (WT) and liver-specific PSAT1 knockout (KO) mice fed a high-fat control (HF-CTRL) or HF-SAAR diet for 6 weeks. **C.** Liver methionine, cysteine, taurine, reduced glutathione (GSH), and oxidized glutathione (GSSG; arbitrary units) in 7-hour fasted mice. **D.** Fractional enrichment (% of total pool) of liver methionine, taurine, GSH, and GSSG following a 30-minute intravenous infusion of [U-^13^C]glucose. **E.** Fractional enrichment (% of total pool) of liver serine, methionine, taurine, GSH, and GSSG at 15 minutes following an i.p. injection of [U-^13^C]serine. n=6-9. *p<0.05, **p<0.01, ***p<0.001, ****p<0.0001.

### 3.5. SAAR reduces glyoxalase cycle intermediates and associated protein PTMs

To further explore the potential connectivity between serine synthesis and GSH availability, we assessed markers of glyoxalase cycle metabolism (Fig. 6A). GSH can be used to generate LGSH in the glyoxalase cycle and recent work identified LGSH-derived lysine lactoylation inhibits PHGDH [42]. Liver glyoxalase cycle enzymes, glyoxalase 1 (GLO1) and hydroxyacylglutathione hydrolase (GLO2), were comparable between genotypes on both diets (Fig. 6B). However, SAAR decreased liver GLO1 protein (Fig. 6B). Consistent with data from our untargeted metabolomic approaches (Fig. 5C), targeted quantification showed lower liver GSH in WT and KO mice fed a HF-SAAR diet compared to the HF-CTRL diet (Fig. 6C). Dietary SAAR also decreased liver LGSH and HTA in both genotypes (Fig 6C). Liver lactate was lower in liver PSAT1 KO mice relative to WT mice fed HF-CTRL (Fig. 6C). SAAR decreased liver lactate in WT, but not liver PSAT1 KO mice (Fig. 6C). PTMs derived from glyoxalase cycle metabolites were broadly suppressed by SAAR. Specifically, lysine lactoylation was decreased in HF-SAAR WT mice compared to WT HF-CTRL (Fig. 6D). In contrast, lysine lactoylation was comparable between liver PSAT1 KO mice fed the HF-CTRL and-SAAR diets (Fig. 6D). Methylglyoxal (MGO)-derived modifications, MG-H1 and CEA, were significantly lower in HF-SAAR-versus HF-CTRL-fed mice regardless of genotype (Fig. 6D). Global lysine acetylation was not changed by diet or genotype (Fig. 6D). Overall, SAAR reduces glyoxalase-cycle intermediates and the protein modifications they support.

**Figure 6.**
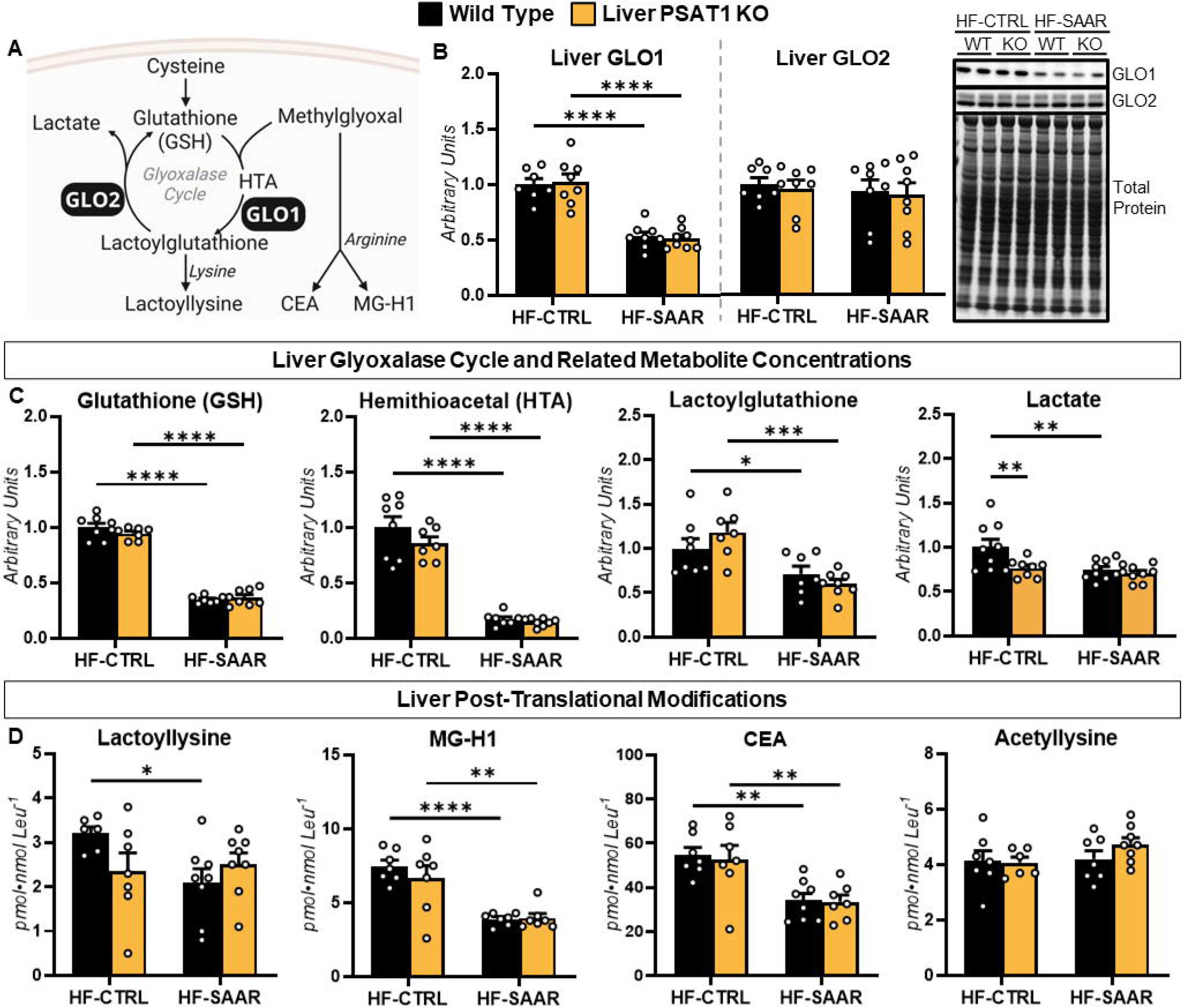
Sulfur amino acid restriction (SAAR) lowers glyoxalase cycle metabolites and associated post-translational modifications (PTMs). **A.** A schematic of select reactions, metabolites, and PTMs related to the glyoxalase cycle. **B.** Liver glyoxalase 1 (GLO1) and hydroxyacylglutathione hydrolase (GLO2) as determined by immunoblotting and representative immunoblots in wild type (WT) and liver-specific PSAT1 knockout (KO) mice fed a high-fat control (HF-CTRL) or HF-SAAR diet for 6 weeks. **C.** Liver glutathione (GSH; arbitrary units), hemithioacetal (HTA; arbitrary units), lactoylglutathione (arbitrary units), and lactate (arbitrary units). **D.** Global PTMs (pmol·nmol Leu^-1^): lactoyllysine, methylglyoxal-derived hydroimidazolone 1 (MG-H1), carboxyethylarginine (CEA), and acetyllysine. n=6-9. *p<0.05, **p<0.01, ****p<0.0001.

### 3.6. SAAR diverts glucose toward asparagine synthesis

Given that loss of hepatic PSAT1 blunted, but did not completely abolish the improvement in glucose tolerance by SAAR, we further examined how SAAR could be influencing glucose metabolism in the liver. On a HF-CTRL diet, liver TCA cycle, anaplerotic, and cataplerotic metabolites were mostly comparable between WT and KO mice (Fig. 7A and B). SAAR increased pyruvate, decreased malate (WT mice only), and elevated aspartate (Fig. 7B). [U-^13^C]glucose tracing showed lower ^13^C-pyruvate, increased ^13^C-aspartate, and significantly elevated ^13^C-asparagine (∼370-fold) labeling in WT and KO mice on the HF-SAAR diet compared to the HF-CTRL diet (Fig. 7C). Interestingly, SAAR lowered the ^13^C-labeling of liver succinate in WT mice (Fig. 7C). This decreased ^13^C-labeling of liver TCA cycle intermediates was more pronounced in PSAT1 KO mice on a HF-SAAR diet as indicated by lower ^13^C-citrate, ^13^C-succinate, and ^13^C-malate (Fig. 7C). Next, we assessed three isotopologue ratios to provide indices of nutrient flux [43–48]. Livers of WT mice fed the HF-SAAR diet had a higher citrate M+3-to-pyruvate M+3 ratio relative to WT mice fed a HF-CTRL and KO mice fed a HF-SAAR diet (Fig. 7D). This suggests that TCA cycle anaplerosis via pyruvate carboxylase (PC) was stimulated by SAAR. The aspartate M+3-to-succinate M+3 and aspartate M+3-to-citrate M+3 ratios were elevated in HF-SAAR mice compared to their HF-CTRL counterparts (Fig. 7D). The increase in these two ratios suggest a preferential partitioning of pyruvate to aspartate compared to flux through the TCA cycle. Consistent with the metabolite and ^13^C-isotope tracing data, liver asparagine synthetase (ASNS) protein was ∼25-fold higher in mice fed a HF-SAAR diet compared to those receiving the HF-CTRL diet (Fig. 7E). Most TCA cycle proteins tested were not altered by diet or genotype (Fig. 7E). The cataplerotic enzyme, mitochondrial glutamic-oxaloacetic transaminase (GOT2), was increased in PSAT1 KO mice fed a HF-SAAR diet compared to both their WT littermates and KO mice fed HF-CTRL (Fig. 7E). These results indicate that SAAR shifts the fate of glucose, via the TCA cycle anaplerosis, towards liver asparagine synthesis.

**Figure 7.**
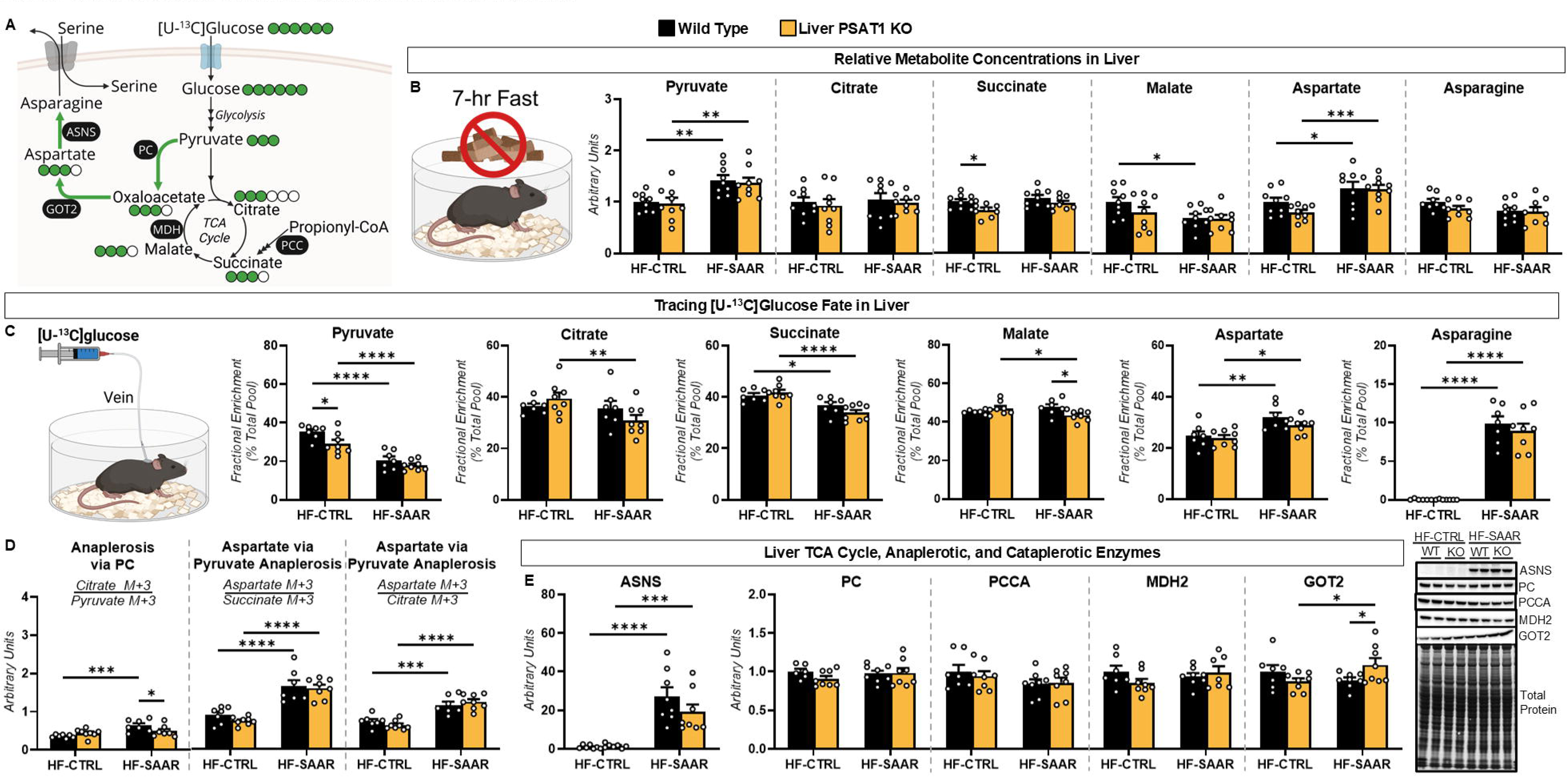
Sulfur amino acid restriction (SAAR) increases glucose partitioning toward asparagine synthesis. **A.** A schematic representation of select reactions and metabolites linking tricarboxylic acid (TCA)-cycle intermediates to asparagine synthesis. **B.** Liver TCA cycle, anaplerotic and cataplerotic metabolites (arbitrary units) in wild type (WT) and liver-specific PSAT1 knockout (KO) mice fed a high-fat control (HF-CTRL) or HF-SAAR diet for 6 weeks. **C.** Fractional enrichment (% of total pool) of liver TCA cycle, anaplerotic and cataplerotic metabolites following a 30-minute intravenous infusion of [U-^13^C]glucose. **D.** Ratios of isotopologues used as indices of anaplerotic and cataplerotic flux. **E.** Liver asparagine synthase (ASNS), pyruvate carboxylase (PC), propionyl-CoA carboxylase subunit A (PCCA), malate dehydrogenase 2 (MDH2), and mitochondrial glutamic-oxaloacetic transaminase (GOT2) as determined by immunoblotting and representative immunoblots. n=6-9. *p<0.05, **p<0.01, ***p<0.001, ****p<0.0001.

### 3.7. SAAR prevents liver lipid accumulation independently of hepatic PSAT1

In addition to glucose control, we investigated the role of hepatic PSAT1 action in liver lipid adaptations to dietary SAAR. After 6 weeks of diet, liver triglycerides were lower in WT mice fed a HF-SAAR diet compared to WT mice on the HF-CTRL diet (Fig. 8A). This dietary effect was not observed in liver PSAT1 KO mice (Fig. 8A). The expression of liver *de novo* lipogenesis enzymes, ATP-citrate lyase (ACLY) and stearoyl-CoA desaturase 1 (SCD1), were suppressed with HF-SAAR feeding in both genotypes (Fig. 8B and D). SAAR decreased FASN protein in WT mice only (Fig. 8B and D). Given that hepatic triglyceride concentrations may also be impacted by changes in lipid catabolism, we next examined mitochondrial respiratory proteins as a marker of potential changes in mitochondrial oxidative metabolism in response to SAAR. As we previously observed [21], SAAR increased liver mitochondrial respiratory proteins (Fig. 8C and D). Specifically, complex I (NADH dehydrogenase (ubiquinone) iron-sulfur protein 4; NDUFS4) and complex V (ATP synthase F1 subunit alpha; ATP5A) were higher in WT mice fed HF-SAAR compared to WT mice fed the HF-CTRL diet. (Fig. 8C and D). Complex IV (mitochondrial encoded cytochrome c oxidase subunit II; MTCO2) protein was elevated in PSAT1 KO mice on the HF-SAAR diet versus KO mice fed the HF-CTRL diet (Fig. 8C and D). An additional mechanism through which SAAR may lower liver triglycerides is the formation of acylglycines (Fig. 8E). Protein expression of liver glycine N-acyltransferase (GLYAT), which catalyzes the conjugation of acyl-CoA to glycine, was similar between groups (Fig. 8F). However, HF-SAAR raised the levels of short-chain acylglycines (acetylglycine, isobutyrylglycine, and isovalerylglycine) in the liver (Fig. 8G).

**Figure 8.**
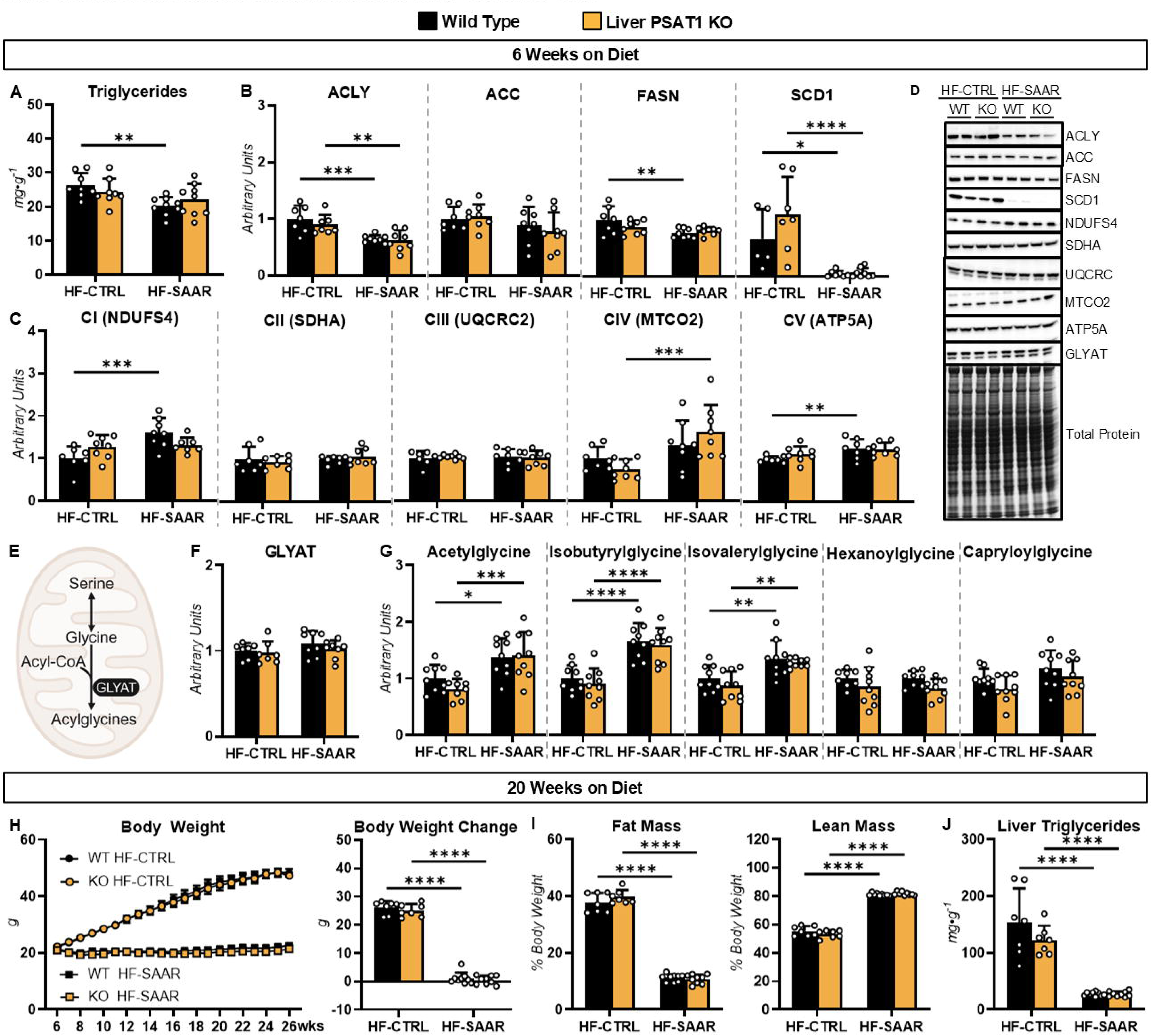
Sulfur amino acid restriction (SAAR) reduces lipid accumulation independently of hepatic PSAT1. Panels A-G are measurements in in wild type (WT) and liver-specific PSAT1 knockout (KO) mice fed a high-fat control (HF-CTRL) or HF-SAAR diet for 6 weeks. Panels H-J are measurements in WT and KO mice fed the HF-CTRL and-SAAR diets for 20 weeks. **A.** Liver triglycerides (mg·g^-1^). **B.** Liver *de novo* lipogenesis enzymes: ATP-citrate lyase (ACLY), acetyl-CoA carboxylase (ACC), fatty acid synthase (FASN), and stearoyl-CoA desaturase 1 (SCD1) as determined by immunoblotting (arbitrary units). **C.** Liver mitochondrial respiratory complexes I-V (CI-V) as determined by immunoblotting. **D.** Representative immunoblots for panels B, C, and F. **E.** A schematic representation of select metabolites and enzymes related to the synthesis of acylglycines. **F.** Liver glycine N-acyltransferase (GLYAT) as determined by immunoblotting. **G.** Liver acylglycines (arbitrary units): acetylglycine, isobutyrylglycine, isovalerylglycine, hexanoylglycine, and capryloylglycine. **H.** Body weight time course (g) and change in body weight (g) over 20 weeks on diet. **I.** Fat mass and lean mass (% body weight). **J.** Liver triglycerides (mg·g^-1^). n=5-9 *p<0.05, **p<0.01, ***p<0.001, ****p<0.0001.

To further test if hepatic PSAT1 action was required for SAAR to lower liver triglycerides, mice were fed the HF-SAAR diet for 20 weeks. Both WT and KO mice fed the HF-SAAR diet had lower body weight, fat mass, and liver triglycerides compared to mice on the HF-CTRL diet (Fig. 8H-J). This suggest that hepatic PSAT1 is not needed, particularly with increasing duration, for SAAR to promotes a decline in liver triglycerides.

## 4. DISCUSSION

Dietary SAAR prevents obesity, dysregulated glucose control, and liver steatosis [1]. Analysis of changes at the level of the transcriptome has identified that the liver is particularly responsive to SAAR [1]. These changes include an increased expression of liver *de novo* serine synthesis pathway enzyme, PSAT1, which is among the most reproducible responses to dietary SAAR [2; 9; 10; 15–23]. Given the interconnectedness of *de novo* serine synthesis with glycolysis and gluconeogenesis pathways, this study tested the hypothesis that hepatic PSAT1 is needed for the liver glucose and lipid adaptations to SAAR. The pertinent findings of this study are: 1) enhanced hepatic *de novo* serine synthesis is required for SAAR to improve glucose tolerance, 2) concurrent with transcriptional activation, a serine synthesis-GSH-lysine lactoylation feedback loop may underlie the increased hepatic serine synthesis in response to SAAR, and 3) hepatic serine synthesis is not necessary for the SAAR-mediated decline in adiposity and liver triglycerides.

### 4.1. Role of hepatic PSAT1 in glucose adaptations to SAAR

Consistent with previous studies using rodents [2–7], we found that dietary SAAR enhances glucose tolerance. Untargeted metabolomics showed this whole-body glucose phenotype was associated with increased liver serine. Exogenous serine supplementation can [49], but does not always [50], improve glucose tolerance. Thus, the SAAR-mediated increase in serine may actively promote glucose tolerance via one or more downstream functions [51]. Our ^13^C-isotope tracing experiments showed elevated partitioning of circulating glucose to hepatic serine following SAAR. Moreover, loss of hepatic PSAT1 impaired glucose tolerance in mice fed the SAAR diet and abolished the increase in liver serine sourced from circulating glucose without completely preventing the rise in liver serine levels. These results suggest that a higher rate of hepatic glucose disposal and the diversion of glucose to the synthesis of serine is, in part, responsible for the SAAR-mediated adaptations in whole-body glucose control.

PSAT1 overexpression in mice enhances both glucose and insulin tolerance [26]. In the experiments presented here, we found that the heightened insulin tolerance following SAAR was not impacted by the loss of hepatic PSAT1. Prior work has reported that the enhanced insulin action in response to PSAT1 overexpression is mediated by decreased expression of tribbles homolog 3 (TRB3), which is a pseudokinase that inhibits insulin signaling by binding to Akt and impairing its activation by phosphorylation [26]. Our studies showed the phosphorylation of liver Akt was elevated similarly in both WT and liver PSAT1 KO mice. Together, these data suggest the elevated liver PSAT1 increases glucose tolerance independent of insulin action. However, future work should assess the role of liver PSAT1 on liver insulin action during hyperinsulinemic-euglycemic clamps combined with isotope tracers. These conditions would allow for assessing liver-specific insulin sensitivity under euglycemic conditions so that the impact of SAAR on extrahepatic glucose flux and concentrations do not mask alterations at the liver.

### 4.2. Potential mechanisms promoting increased hepatic de novo serine synthesis from glucose

Work by independent groups indicate that SAAs negatively regulate serine synthesis. Cysteine suppresses the expression of PHGDH by decreasing mRNA stability [52]. In addition to post-transcriptional regulation, mechanisms at the level of transcription may also contribute to the enhanced serine synthesis by SAAR. Deletion of the transcription factor, nuclear factor erythroid 2-related factor 2 (NRF2), in the livers of mice blunted but did not abolish the increase in PSAT1 mRNA [16]. These mechanisms of action are consistent with the elevated liver PHGDH and PSAT1 mRNA and protein in response to SAAR. Serine is a precursor for cysteine, which is generated via transsulfuration wherein the sulfur from methionine is transferred to serine [41]. It may be that increased cysteine impedes the synthesis of its precursor, serine, to control SAA levels. Conversely, when cysteine is low, serine synthesis is stimulated to promote SAA availability. Ultimately, these findings suggest that SAAR-mediated increase in glucose tolerance is the result of liver responses to maintain SAA homeostasis via serine synthesis.

A prominent fate of methionine and cysteine, particularly under conditions of low SAA levels, is the synthesis of GSH [53–56]. Aligning with prior studies [23], we observed liver GSH was decreased by SAAR. However, our [U-^13^C]glucose and [U-^13^C]serine isotope tracing experiments showed that ^13^C-labeling of liver GSH and GSSG was elevated in mice fed the SAAR diet. While the [U-^13^C]serine tracing showed SAAR increased labeling of other fates of cysteine such as taurine, loss of hepatic PSAT1 uniquely attenuated the increased liver ^13^C-GSH and ^13^C-GSSG from [U-^13^C]glucose. These results indicate that serine synthesized *de novo* from glucose is important to support GSH provision (via cysteine) during SAAR.

Beyond its antioxidant properties, GSH is a metabolite within the glyoxalase cycle. This highly conserved metabolic cycle comprised of two enzymes, GLO1 and GLO2, uses GSH to catabolize MGO to lactate through the intermediate, LGSH [57]. LGSH is an acyl-donor in a non-enzymatic reaction with free lysine residues, yielding the PTM, lysine lactoylation [35; 58]. This is notable because PHGDH and PSAT1 are heavily lactoylated and LGSH inhibits PHGDH activity [35; 42]. Our results showed that SAAR lowered liver LGSH and lysine lactoylation. Together, our data and the work of others suggest that SAAR may increase serine synthesis by preventing the inhibition of PHGDH and/or PSAT1 by LGSH-derived lysine lactoylation. Consistent with GSH regulating serine synthesis via post-translational mechanisms, the pharmacologic inhibition of GSH synthesis using buthionine sulfoximine increased liver serine without altering NRF2 or PHGDH expression [22]. While it remains to be fully tested, SAAR may promote glucose tolerance through a hepatic serine synthesis-GSH-lysine lactoylation feedback loop in which low SAA decreases liver GSH, lowers lysine lactoylation, and thereby prevents inhibition of *de novo* serine synthesis from glucose.

### 4.3. SAAR shifts the fate of glucose towards liver asparagine

In addition to *de novo* serine synthesis, this study demonstrates that SAAR stimulates glucose partitioning towards asparagine synthesis. Liver asparagine concentration was similar between mice fed the HF-CTRL and SAAR diets. However, ^13^C-labeling of liver asparagine from [U-^13^C]glucose increased markedly by ∼370-fold in SAAR-fed mice. We also assessed indices of nutrient flux [43–48]. Isotopologue ratios from our [U-^13^C]glucose tracing experiments are consistent with SAAR increasing TCA cycle anaplerosis through PC (i.e., the generation of OAA from pyruvate) and the resulting OAA being preferentially directed to asparagine rather than flux through the TCA cycle. In agreement with this, Xin et al.[59] show that SAAR increases liver TCA cycle anaplerosis as evidenced by higher enrichment of TCA cycle metabolites following the administration of [U-¹³C]glucose and [U-¹³C,¹⁵N]glutamine. Herein, we show the SAAR-induced shift in glucose and TCA cycle intermediates to asparagine was associated with a ∼25-fold increase in liver ASNS, which generates asparagine from aspartate. SAAR did not alter the expression of other anaplerotic and cataplerotic enzymes (i.e., PC, MDH2, PCCA, and GOT2) linked to asparagine synthesis. Interestingly, these aforementioned enzymes have been reported to be modified by lysine lactoylation [35]. Thus, further research is warranted to determine if the SAAR-driven reduction in lactoylation prevents the inhibition of these enzymes and favors the diversion of glucose to asparagine. Future studies should also test whether the stimulation glucose partitioning to asparagine is an additional means through which SAAR increases glucose tolerance. The shift in glucose fate to asparagine during SAAR could be an additional means of enhancing serine availability. Intracellular asparagine exchanges with extracellular serine to promote serine uptake [60]. Liver PSAT1 KO mice receiving a HF-SAAR diet showed higher liver serine levels compared to KO mice fed a HF-CTRL diet. This was accompanied by a trend towards greater ^13^C-labeling of liver serine [U-^13^C]glucose in KO mice fed a HF-SAAR diet compared to KO mice fed a HF-CTRL diet. To further explore if the liver was acquiring this serine from extrahepatic sources, we administered [U-^13^C]serine to mice. Liver ^13^C-serine labeling was similar between mice fed a HF-CTRL and HF-SAAR diet. Given that the liver serine pool size is larger in mice fed a HF-SAAR diet, a greater amount of exogenous ^13^C-serine would need to be taken up by the liver to achieve a similar proportion of ^13^C-serine labeling in livers of mice fed a HF-SAAR diet. Thus, our data aligns with the hypothesis that SAAR increases asparagine synthesis from glucose to promote liver serine uptake.

### 4.4. Role of hepatic PSAT1 in lipid adaptations to SAAR

Our findings agree with prior studies reporting that SAAR lowers adiposity and liver triglycerides [2; 5; 9; 18; 21; 59]. In the current study, the decline in liver triglycerides following SAAR was associated with decreased *de novo* lipogenesis enzymes (ACLY, FASN, and SCD-1). There was a modest increase in mitochondrial respiratory chain proteins (complexes I, IV, and V) in response to SAAR. These results are consistent with our prior work showing dietary SAAR increases the expression of select mitochondrial proteins without enhancing mitochondrial respiration [21]. While terminal oxidation may not be enhanced by SAAR [21], other means of lipid disposal could contribute to lower triglycerides. Studies using ^14^C-palmitate in liver homogenates have determined that SAAR increases liver ꞵ-oxidation [61]. We identified short-chain acylglycines (acetylglycine, isobutyrylglycine, and isovalerylglycine) were increased. These acylglycines are generated in the mitochondria by conjugating acyl-CoAs to glycine in a reaction mediated by GLYAT [62]. Thus, the SAAR-mediated rise in glycine may facilitate acylglycine synthesis and, consequently, promote the disposal of fatty acids during SAAR. Importantly, SAAR elicited the molecular responses in lipid metabolism and lowered liver triglycerides similarly in WT and PSAT1 KO mice. Overall, the prevailing signature under SAAR is a coordinated decrease in *de novo* lipogenesis and stimulation of acyl-CoA disposal that is independent of PSAT1 action.

In summary, hepatic PSAT1 is necessary for adaptations to SAAR involving serine synthesis that allow improvements in glucose tolerance to be fully realized. In contrast, PSAT1 is dispensable for the adaptations in lipid accretion and disposal pathways that contribute to the SAAR-mediated decline in liver lipids. Thus, the serine synthesis pathway is critical to the glucose, but not lipid, adaptations to SAAR. These studies provide a mechanistic foundation for how SAAR impacts the liver to prevent dysregulated glucose control.

## ACKNOWLEDGEMENTS

The authors acknowledge the technical assistance provided by Ruth Pfeiffer. Figure schematics were created using BioRender.com.

## CRediT AUTHORSHIP CONTRIBUTION STATEMENT

A.F.O.: Writing – original draft, Visualization, Validation, Methodology, Investigation, Formal analysis, Data curation, Conceptualization, and Project administration C.V., F.I.R., K.M.A., A.M.P., and A.B.N.: Investigation, and Writing - Review & Editing; P.A.C., and J.J.G.: Methodology, Resources, and Writing - Review & Editing; S.C.H.: Resources, and Writing - Review & Editing; C.C.H.: Writing – review & editing, Visualization, Validation, Supervision, Resources, Methodology, Investigation, Funding acquisition, Dara curation, Conceptualization, and Project administration.

---

**Supplementary Figure 1. Sulfur Amino Acid Restriction (SAAR) impacts systemic glucose control and hepatic glucose partitioning in C57BL/6J mice independent of insulin levels. A.** A schematic representation of the study design. C57BL/6J male mice were fed a low-fat control (LF-CTRL), high-fat control (HF-CTRL), or HF-SAAR diet for 6 weeks starting at 6 weeks of age. A subset of mice fed the HF-SAAR diet had a subcutaneous insulin pellet (HF-SAAR + Insulin) implanted at 7 weeks of age for sustained release of insulin until 12 weeks of age. **B.** Body weight time course (g), body weight change over the 6 weeks on diet (g), fat mass (% of body weight), and lean mass (% of body weight). **C.** Liver triglycerides (mg·g^-1^) and liver weight (g). **D.** Liver glycogen (mg·g^-1^)**. E.** Blood glucose (mg·dL^-1^). **F.** Glucose excursion and area under the curve (AUC) during a glucose tolerance test. **G.** Glucose excursion and AUC during an insulin tolerance test. **H.** Plasma insulin (pmol·L^-1^). **I.** Liver methionine, homocysteine, cysteine, and taurine (arbitrary units). **J.** Liver phosphoglycerate dehydrogenase (PHGDH) and phosphoserine aminotransferase 1 (PSAT1) as determined by immunoblotting and representative immunoblots. **K.** Liver serine and glycine (arbitrary units). n=3-17. ^#^p<0.05 by *t*-test. *p<0.05, **p<0.01, ***p<0.001, ****p<0.0001 by ANOVA.

