## Supplementary figures and images for "Dietary sulfur amino acid restriction improves glucose homeostasis through hepatic *de novo* serine synthesis"

### Supplementary Figure 1

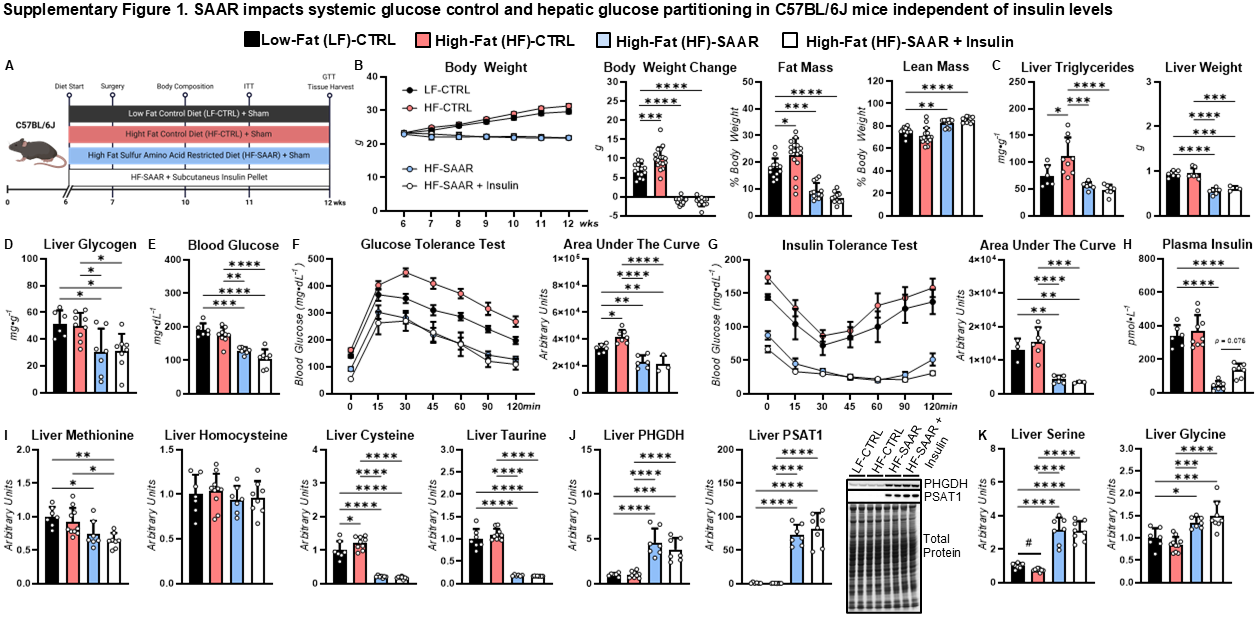
