## Supplementary Table 1 for "Dietary sulfur amino acid restriction improves glucose homeostasis through hepatic *de novo* serine synthesis"

| Diet | Low-Fat Control | | High-Fat Control | | High-Fat SAAR | |
|---|---|---|---|---|---|---|
| **Supplier** | Research Diets, Inc. | | Research Diets, Inc. | | Research Diets, Inc. | |
| **Product #** | A11051302B | | A11051306 | | A11051305 | |
| **Unit** | g/100g diet | kcal% | g/100g diet | kcal% | g/100g diet | kcal% |
| **Carbohydrate** | 74 | 76 | 35 | 27 | 35 | 27 |
| **Fat** | 4 | 10 | 35 | 60 | 35 | 60 |
| **Protein** | 13 | 14 | 18 | 14 | 18 | 14 |
| **Unit (AA)** | g | kcal | g | kcal | g | kcal |
| **L-Arginine** | 11.2 | 45 | 11.2 | 45 | 11.2 | 45 |
| **L-Histidine** | 3.3 | 13 | 3.3 | 13 | 3.3 | 13 |
| **L-Isoleucine** | 8.2 | 33 | 8.2 | 33 | 8.2 | 33 |
| **L-Leucine** | 11.1 | 44 | 11.1 | 44 | 11.1 | 44 |
| **L-Lysine** | 14.4 | 58 | 14.4 | 58 | 14.4 | 58 |
| **DL-Methionine** | 8.6 | 34 | 8.6 | 34 | 1.2 | 5 |
| **L-Phenylalanine** | 11.6 | 46 | 11.6 | 46 | 11.6 | 46 |
| **L-Threonine** | 8.2 | 33 | 8.2 | 33 | 8.2 | 33 |
| **L-Tryptophan** | 1.8 | 7 | 1.8 | 7 | 1.8 | 7 |
| **L-Valine** | 8.2 | 33 | 8.2 | 33 | 8.2 | 33 |
| **L-Glutamic acid** | 27 | 108 | 27 | 108 | 34.4 | 138 |
| **L-Glycine** | 23.3 | 93 | 23.3 | 93 | 23.3 | 93 |
| **Corn starch** | 424.5 | 1698 | 0 | 0 | 0 | 0 |
| **Maltodextrin** | 125 | 500 | 56.8 | 227 | 56.8 | 227 |
| **Dextrose** | 50 | 200 | 50 | 200 | 50 | 200 |
| **Sucrose** | 150 | 600 | 150 | 600 | 150 | 600 |
| **Cellulose** | 50 | 0 | 50 | 0 | 50 | 0 |
| **Lard** | 0 | 0 | 219 | 1971 | 219 | 1971 |
| **Corn oil** | 46 | 414 | 46 | 414 | 46 | 414 |
| **Mineral mix S10001** | 35 | 0 | 35 | 0 | 35 | 0 |
| **Vitamin mix V10001** | 10 | 40 | 10 | 40 | 10 | 40 |
| **Choline bitartrate** | 2 | 0 | 2 | 0 | 2 | 0 |
| **Total** | 1029.45 | 4000 | 755.75 | 4000 | 755.75 | 4000 |

**Supplementary Table 1. Dietary composition of low-fat control diet, high-fat control, and high-fat sulfur amino acid restricted (SAAR) diets.**
