## Supplementary Table 2 for "Dietary sulfur amino acid restriction improves glucose homeostasis through hepatic *de novo* serine synthesis"

| Antibody | Supplier | Catalog Number | RRID | Dilution |
|---|---|---|---|---|
| Acetyl-CoA carboxylase | Cell Signaling Technologies | 3662 | RRID:AB_2219400 | 1:1000 |
| Phospho-Akt (Ser473) | Cell Signaling Technologies | 9271 | RRID:AB_329825 | 1:1000 |
| Akt | Cell Signaling Technologies | 9272 | RRID:AB_329827 | 1:1000 |
| Asparagine Synthetase | Cell Signaling Technologies | 20843 | N/A | 1:1000 |
| ATP-citrate lyase | Cell Signaling Technologies | 4332 | RRID:AB_2223744 | 1:1000 |
| Cystathionine β-synthase | Proteintech | 14787-1-AP | RRID:AB_2070970 | 1:2000 |
| Cysteine dioxygenase, type I | Proteintech | 12589-1-AP | RRID:AB_10638145 | 1:1000 |
| Cystathionine γ-lyase | Proteintech | 12217-1-AP | RRID:AB_2087497 | 1:1000 |
| Fatty acid synthase | Cell Signaling Technologies | 3180 | RRID:AB_2100796 | 1:1000 |
| Glutamate-cysteine ligase | Proteintech | 12601-1-AP | RRID:AB_2278734 | 1:1000 |
| Glutamic-oxaloacetic transaminase 2 | Proteintech | 14800-1-AP | RRID:AB_2247898 | 1:5000 |
| Glutathione synthetase | Proteintech | 15712-1-AP | RRID:AB_2878171 | 1:1000 |
| Glycine N-acyltransferase | Proteintech | 10900-1-AP | RRID:AB_2232373 | 1:1000 |
| Phospho-glycogen phosphorylase (Ser15) | Abcam | ab227043 | RRID:AB_3674858 | 1:1000 |
| Glycogen phosphorylase | Proteintech | 15851-1-AP | RRID:AB_2175014 | 1:1000 |
| Phospho-glycogen synthase (Ser641) | Cell Signaling Technologies | 3891 | RRID:AB_2116390 | 1:1000 |
| Glycogen synthase 2 | Proteintech | 22371-1-AP | RRID:AB_2879091 | 1:1000 |
| Glyoxalase I | Proteintech | 15140-1-AP | RRID:AB_2109890 | 1:1000 |
| Hydroxyacylglutathione hydrolase | Proteintech | 17196-1-AP | RRID:AB_2878361 | 1:1000 |
| Malate dehydrogenase 2 | Proteintech | 15462-1-AP | RRID:AB_2878143 | 1:1000 |
| Mitochondrial encoded cytochrome c oxidase II | Proteintech | 55070-1-AP | RRID:AB_10859832 | 1:1000 |
| NADH dehydrogenase (ubiquinone) iron-sulfur protein 4 | Abcam | ab139178 | RRID:AB_2922810 | 1:1000 |
| Phosphoglycerate dehydrogenase | Cell Signaling Technologies | 13428 | RRID:AB_2750870 | 1:1000 |
| phosphoserine aminotransferase 1 | Proteintech | 10501-1-AP | RRID:AB_2172597 | 1:1000 |
| Propionyl coenzyme A carboxylase α | Proteintech | 21988-1-AP | RRID:AB_2878963 | 1:1000 |
| Pyruvate Carboxylase | Proteintech | 16588-1-AP | RRID:AB_1851513 | 1:5000 |
| Serine hydroxymethyltransferase 2 | Proteintech | 11099-1-AP | RRID:AB_2188452 | 1:1000 |
| Stearoyl-CoA desaturase 1 | Cell Signaling Technologies | 2794 | RRID:AB_2183099 | 1:1000 |
| Succinate dehydrogenase subunit A | Cell Signaling Technologies | 5839 | RRID:AB_10707493 | 1:1000 |
| Total OXPHOS Rodent WB Antibody Cocktail | Abcam | ab110413 | RRID:AB_2629281 | 1:1000 |
| Anti-rabbit IgG | Cell Signaling Technologies | 7074 | RRID:AB_2099233 | 1:5000 |
| Anti-mouse IgG | Cell Signaling Technologies | 7076 | RRID:AB_330924 | 1:5000 |

**Supplementary Table 2. List of antibodies.**
