## Supplementary Table 3 for "Dietary sulfur amino acid restriction improves glucose homeostasis through hepatic *de novo* serine synthesis"

| Diet | Low-Fat Control | | Low-Fat SAAR | |
|---|---|---|---|---|
| Supplier | Dyets Inc. | | Dyets Inc. | |
| Product # | 510072 | | 510071 | |
| Unit | g/100g diet | kcal% | g/100g diet | kcal% |
| Carbohydrate | 68.25 | 68 | 68.25 | 68 |
| Fat | 8 | 18 | 8 | 18 |
| Protein | 14.05 | 14 | 14.05 | 14 |
| Unit (AA) | g | kcal | g | kcal |
| L-Arginine | 11.2 | 45 | 11.2 | 45 |
| L-Histidine | 3.3 | 13 | 3.3 | 13 |
| L-Isoleucine | 8.2 | 33 | 8.2 | 33 |
| L-Leucine | 11.1 | 44 | 11.1 | 44 |
| L-Lysine | 18 | 72 | 18 | 72 |
| DL-Methionine | 8.6 | 34 | 1.72 | 7 |
| L-Phenylalanine | 11.6 | 46 | 11.6 | 46 |
| L-Threonine | 8.2 | 33 | 8.2 | 33 |
| L-Tryptophan | 1.8 | 7 | 1.8 | 7 |
| L-Valine | 8.2 | 33 | 8.2 | 33 |
| L-Glutamic acid | 27 | 108 | 33.88 | 136 |
| (Glycine | 23.3 | 93 | 23.3 | 93 |
| Corn starch | 432.5 | 1557 | 432.5 | 1557 |
| Maltodextrin | 0 | 0 | 0 | 0 |
| Dyetrose | 50 | 190 | 50 | 190 |
| Dextrose | 200 | 800 | 200 | 800 |
| Sucrose | 0 | 0 | 0 | 0 |
| Cellulose | 50 | 0 | 50 | 0 |
| Lard | 0 | 0 | 0 | 0 |
| Corn oil | 80 | 720 | 80 | 720 |
| Salt mix #200000 | 35 | 16.45 | 35 | 16.45 |
| Vitamin mix #300050 | 10 | 38.7 | 10 | 39.2 |
| Choline bitartrate | 2 | 0 | 2 | 0 |
| Total | 1000 | 3884.15 | 1000 | 3884.15 |

**Supplementary Table 3. Dietary composition of low-fat control diet and Low-fat sulfur amino acid restricted (SAAR) diets used in Figure 2F-H.**
